# Recombinant and synthetic affibodies function comparably for modulating protein release

**DOI:** 10.1101/2024.05.11.593708

**Authors:** Jonathan Dorogin, Morrhyssey A. Benz, Cameron J. Moore, Danielle S. W. Benoit, Marian H. Hettiaratchi

## Abstract

**Purpose:** Affibodies are a class of versatile affinity proteins with a wide variety of therapeutic applications, ranging from guided contrast agents for imaging to cell-targeting therapeutics. We have identified several affibodies specific to bone morphogenetic protein-2 (BMP-2) with a range of binding affinities and demonstrated the ability to tune release rate of BMP-2 from affibody-conjugated poly(ethylene glycol) (PEG) hydrogels based on affibody affinity strength. In this work, we compare the purity, structure, and activity of recombinant, bacterially-expressed BMP-2-specific affibodies with affibodies synthesized via solid-phase peptide synthesis.

**Methods:** High- and low-affinity BMP-2-specific affibodies were recombinantly expressed using BL21(DE3) *E. coli* and chemically synthesized using microwave-assisted solid-phase peptide synthesis with Fmoc-Gly-Wang resin. The secondary structures of the affibodies and dissociation constants of affibody-BMP-2 binding were characterized by circular dichroism and biolayer interferometry, respectively. Endotoxin levels were measured using chromogenic limulus amebocyte lysate (LAL) assays. Affibody-conjugated PEG-mal hydrogels were fabricated and loaded with BMP-2 to evaluate hydrogel capacity for controlled release, quantified by enzyme-linked immunosorbent assays (ELISA).

**Results:** Synthetic and recombinant affibodies were determined to be α-helical by circular dichroism. The synthetic high- and low-affinity BMP-2-specific affibodies demonstrated comparable BMP-2-binding dissociation constants to their recombinant counterparts. Recombinant affibodies retained some endotoxins after purification, while endotoxins were not detected in the synthetic affibodies. High-affinity affibody-conjugated hydrogels reduced cumulative BMP-2 release compared to the low-affinity affibody-conjugated hydrogels and hydrogels without affibodies.

**Conclusions:** Synthetic affibodies demonstrate comparable structure and function to recombinant affibodies, while reducing endotoxin contamination and increasing product yield, indicating that solid-phase peptide synthesis is a viable method of producing affibodies for controlled protein release and other applications.

## Introduction

Affibodies are a class of versatile antibody-mimetic proteins that can be engineered to bind with high specificity and tunable affinity to various targets, enabling their use in numerous biomedical applications, including contrast agents for medical imaging and affinity-mediated protein delivery vehicles.^1–4^ Recently, we identified several affibodies that specifically bind to bone morphogenetic protein-2 (BMP-2) with a range of affinities and used these affibodies to tune the release of BMP-2 from affibody-conjugated poly(ethylene glycol) hydrogels based on the strength of the affibody-BMP-2 affinity interaction.^5^ In this work, high-affinity and low-affinity BMP-2 affibodies were expressed recombinantly in BL21(DE3) *E. coli*. However, bacterial protein expression can result in contamination with endotoxin, a bacterial membrane component, leading to inflammation and activation of the innate immune system in vivo.^6,7^ Thus, reducing endotoxin levels in affibodies could improve their utility in biomedical applications.

Common approaches to reducing endotoxin contamination include post-processing recombinant proteins with endotoxin removal columns^8–10^ and phase separation methods,^8,11^ using low endotoxin bacterial expression systems and mammalian cell lines to minimize levels of endotoxin expressed,^12–15^ and solid-phase protein synthesis (SPPS).^16^ While recombinant protein expression is optimized for the expression system (i.e., BL21 E. coli) to ensure protein expression and proper folding, the yield can be quite low. Conversely, the higher mass yields associated with SPPS do not guarantee proper protein folding and require multiple purification steps. Further, SPPS synthesis efficiency decreases with larger peptide and protein sequences.^17,18^ As a result, recombinant and synthetic proteins functions may differ slightly. In this work, we compare the yield, endotoxin levels, secondary structure, and affinity binding interactions of recombinant and synthetic affibodies to determine whether synthetic BMP-2-specific affibodies can substitute recombinant affibodies for controlled BMP-2 delivery.

## Methods and Results

Unless otherwise specified, all chemicals were purchased from Fisher Scientific.

### Bacterial expression and purification of recombinant affibodies

Recombinant affibodies were expressed in *E. coli* and purified as previously described.^5^ Briefly, pET28b+ vectors for isopropyl β-D-1-thiogalactopyranoside (IPTG (Thermo Scientific))-inducible protein expression were constructed with high- or low-affinity BMP-2-specific affibody sequences modified with a C-terminal hexahistidine tag for immobilized metal affinity chromatography and cysteine for bioconjugation. Plasmids were transformed into BL21 *E. coli*, and sequences were verified using Sanger sequencing (Azenta Life Sciences). *E. coli* were expanded in 20 mL of Lysogeny Broth overnight at 37 °C, and then further expanded in 1.8 L of Terrific Broth (Research Products International) at 37 °C with air sparging in a bioreactor (LEX-10, Epiphyte3) for 3-4 hours until the optical density of the culture at 600 nm reached approximately 0.8-1.2 absorbance units. Protein expression was induced with IPTG at 18 °C for 20 hours. The cells were lysed by sonication in 50 mM Tris buffer pH 7.5 supplemented with 500 mM NaCl, 5 mM imidazole (Sigma Aldrich), and 75 mg of tris(2-carboxyethyl)phosphine (TCEP (GoldBio)) to prevent oxidation of the cysteines and then centrifuged (13,000 RCF for 30 minutes). The supernatant was mixed with cobalt-nitrilotriacetic acid (Co-NTA (GoldBio)) beads for 1 hour and washed with 100 mL of 50 mM tris buffer containing 500 mM NaCl and 30 mM imidazole. The affibodies were eluted into 50 mM tris buffer pH 7.5 containing 500 mM NaCl and 250 mM imidazole and buffer-exchanged into phosphate-buffered saline (PBS). The affibodies were further purified by size exclusion chromatography (BioRad NGC with Enrich SEC 70, 10/300 mm column), concentrated to 0.5-2 mg/mL, and stored frozen at −20 °C.

### Synthesis and purification of synthetic affibodies

Synthetic affibodies modified with a penultimate cysteine and a C-terminal glycine were prepared by SPPS on Fmoc-Gly-Wang resin (CEM Corporation) (0.602 mmol/g loading) using a CEM Liberty Blue 2.0 microwave peptide synthesizer (CEM Corporation). 0.2 M amino acid (CEM Corporation), 10% v/v pyrrolidine (Sigma Aldrich) deprotection, 1 M N,N’-diisopropylcarbodiimide (DIC) (Oakwood Chemical), and 1 M Oxyma (CEM Corporation) solutions were prepared in DMF. Affibodies were cleaved and deprotected by vacuum-filtering the resin, washing with dichloromethane twice, and resuspending in the cleavage cocktail (1:1:1:1:36 ratios of ddH_2_O, triisopropyl silane (Oakwood Chemical), 2,2’- (Ethylenedioxy)diethanethiol (XX), thioanisole (XX), trifluoroacetic acid (TFA) (Oakwood Chemical)) for 40 minutes at 42 °C^19^. The solution was vacuum filtered and the filtrate was purified via precipitation and three rounds of centrifugation (1200 RCF for 5 minutes) using −20 °C diethyl ether. The resultant slurry was dried overnight in a vacuum desiccator and transferred to −20 °C for storage. The crude products were purified by high-performance liquid chromatography (HPLC; CEM Prodigy using 19 x 150 mm C18 Waters column) by running a 15-75% acetonitrile gradient in ddH_2_O with 0.1% v/v TFA. The purified products were lyophilized and stored at −20 °C.

### Characterization of synthetic and recombinant affibodies

Affibody molecular weights were confirmed using matrix-assisted laser desorption/ionization time of flight (MALDI-TOF) mass spectrometry (Bruker Smart LS). A matrix solution of 10 mg/mL α-cyano-4-hydroxycinnamic acid (Sigma Aldrich) in 0.1% v/v TFA and 50% v/v acetonitrile in ddH_2_O was prepared, and equal parts matrix and sample in 3% v/v acetonitrile in ddH_2_O were deposited on a MALDI target plate.

The secondary structure of the affibodies was measured using circular dichroism (CD) spectrophotometry (Jasco J-815). Affibodies were reconstituted in 10 mM tris buffer, pH 8 and loaded into a quartz cuvette (Starna Cell Corp.) with a 1 mm path length. Circular dichroism spectra were obtained over 190-250 nm and normalized to mean residue molar ellipticity.^20–22^

The kinetics of BMP-2-affibody binding were measured by biolayer interferometry (BLI; Gator Bio). Streptavidin-coated probes were loaded with 25 nM of biotinylated BMP-2 (Medtronic, biotinylated in-house using EZ Link NHS-Biotin (Thermo Fisher) per manufacturer’s protocols)) in PBS with 0.05% Tween 80 (PBST) and then allowed to associate to and dissociate from 31.25-125 mM of affibody in PBST for 60 seconds each. The dissociation constants (K_D_), on-rate constants (*k*_*on*_), and off-rate constants (*k*_*off*_) were calculated for each affibody-BMP-2 binding pair by curve fitting the wavelength shifts using a 1:1 Langmuir isotherm binding model.

### Endotoxin removal and quantification of synthetic and recombinant affibodies

Pierce High Capacity Endotoxin Removal Spin Columns (Thermo Fisher Scientific) were used to remove lipopolysaccharides (LPS), the prominent endotoxin contaminant, from the affibodies. Columns were regenerated and equilibrated using Endotoxin-Free Dulbecco’s PBS (Millipore Sigma) according to the manufacturer’s instructions. Endotoxin levels in each synthetic and recombinant affibody, before and after endotoxin removal, were quantified using the ToxinSensor Chromogenic LAL endotoxin assay kit (GenScript) according to the manufacturer’s instructions. The endotoxin concentration in each sample was determined using a standard curve ranging from 0.1 to 1 endotoxin units (EU)/mL constructed from known amounts of *E. coli* endotoxin, wherein 1 EU corresponds to approximately 0.1-0.2 ng of endotoxin.

### BMP-2 release from hydrogels conjugated with synthetic and recombinant affibodies

Affibody-conjugated PEG-mal hydrogels were synthesized as previously described.^5,23^ Briefly, PEG-maleimide (Laysan Bio) and affibodies (recombinant high-affinity, recombinant low-affinity, synthetic high-affinity, or synthetic low-affinity) were each dissolved in PBS pH 6.9 and rotated for 30 minutes at room temperature to form PEG-affibody intermediates. 70 µL aliquots of intermediate solutions were transferred into 2 mL microcentrifuge tubes, mixed with 30 µL dithiothreitol (DTT) (GoldBio) dissolved in PBS pH 6.9, and allowed to crosslink for 30 minutes at room temperature to form 100 µL hydrogels with 2 nmol of BMP-2-specific affibodies. Unreacted DTT was removed from the hydrogels by washing with fresh PBS 3 times over 36 hours.

Hydrogels were loaded with 100 ng of BMP-2 (3.85 pmol; 500:1 affibody-to-BMP-2 molar ratio) for 3 hours at 4 °C. The remaining loading solution was removed, and BMP-2 concentration was measured using ELISA to calculate initial BMP-2 encapsulation. Hydrogels were submerged in 900 μL of 0.1% w/v bovine serum albumin (BSA) in PBS. BMP-2 release was measured over 2 weeks by collecting 200 μL aliquots of the supernatant periodically and measuring BMP-2 concentration using ELISA. To compare release rates, the slopes of the linear portion of each hydrogel release profile, corresponding to the Fickian diffusion model, were calculated and compared, as previously described.^4,5^

## Results

### Recombinant and synthetic affibodies display similar characteristics

Recombinant expression of BMP-2-specific affibodies in *E. coli* resulted in a yield of approximately 20 mg of purified affibodies from a single 1.8 L culture. SPPS production of affibodies resulted in a yield of approximately 100 mg of purified affibodies from a single 0.1 mmol reaction. MALDI was used to determine the molecular weights of the affibodies. All affibodies displayed single peaks at the expected molecular weights, indicating high purity. The recombinant high-affinity affibody displayed a narrow peak at 7277 Da (−34 Da shift associated with dehydroalanine modifications),^24^ and low-affinity affibody displayed a broader peak at 7627 Da (+213 Da shift associated with myristoilation^25^) (Figure 1A). We have previously reported the presence of these adducts in recombinantly expressed BMP-2-specific affibodies.^5^ The synthetic affibodies both displayed narrow peaks, with the high-affinity affibody displaying a peak at 6544 Da (+2 Da shift associated with reduction of cystine^26^) and the low-affinity affibody displaying a peak at 6650 Da (+2 Da shift associated with reduction of cystine^26^) (Figure 1B). The lower molecular weights of the synthetic affibodies compared to the recombinant affibodies were due to the absence of the hexahistidine tag used in recombinant protein purification.

**Figure 1.**
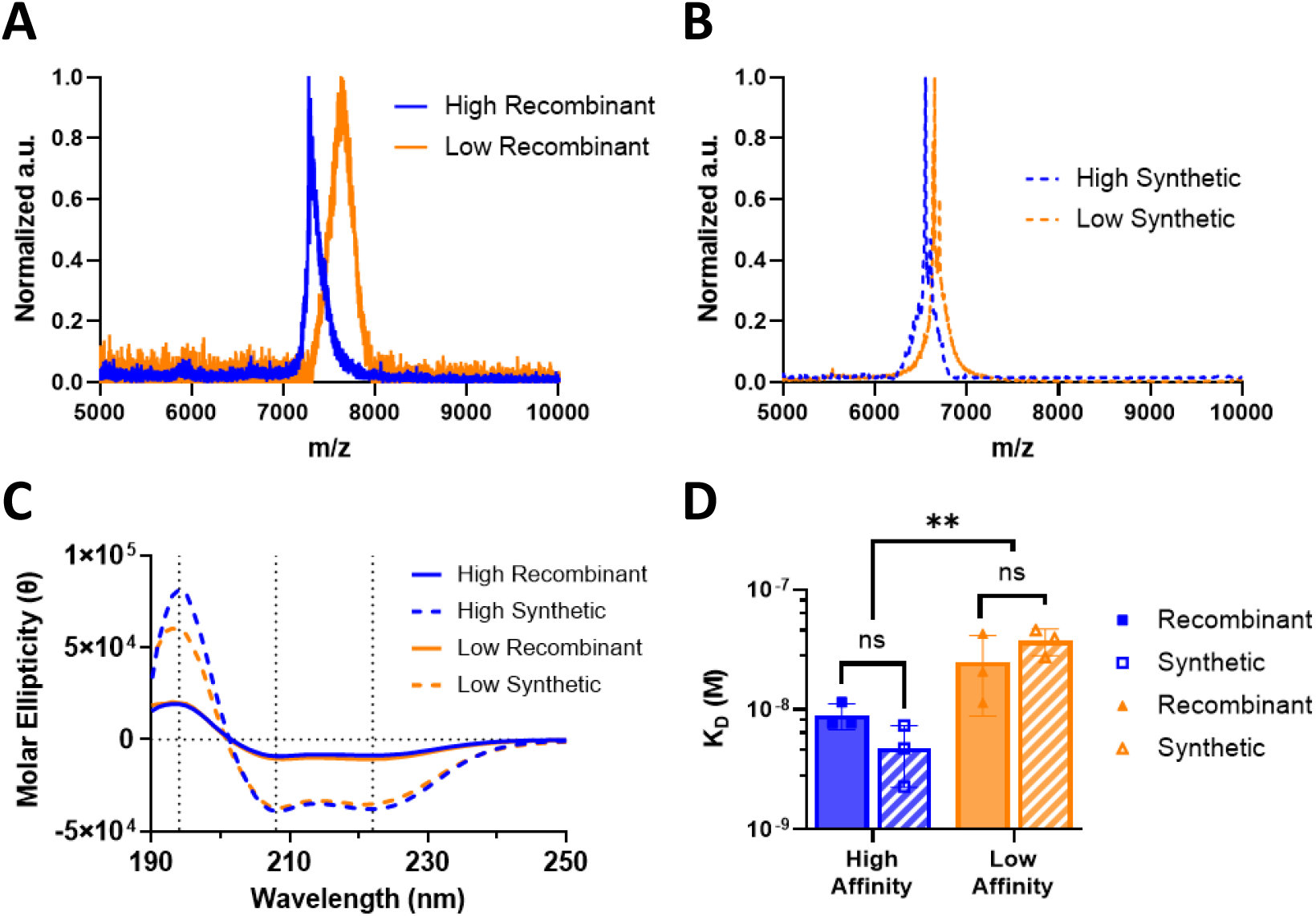
Properties of recombinant and synthetic affibodies. A,B) MALDI spectra of (A) recombinant and (B) synthetic affibodies. A single prominent peak was present in each sample at the expected masses. C) Circular dichroism spectra of affibodies, depicting α-helical secondary structures. D) Equilibrium dissociation constants (K_D_) of affibodies measured by biolayer interferometry. Statistical significance was determined using two-way ANOVA with Tukey post-hoc test. n=3; ns – not significant, ** p<0.01 as indicated.

All affibodies displayed characteristic α-helical CD spectra with troughs at 222 nm and 208 nm, and a peak at 193 nm (Figure 1C).^22^ The increase in molar ellipticity of the synthetic affibodies compared to the recombinant affibodies could be attributed to either the absence of the hexahistidine tag in the synthetic product or the presence of LPS in the recombinant product, as hexahistidine tags do not typically contribute to the α-helicity and reduce the mean residue secondary structure,^27^ while LPS can affect protein folding.^11,28,29^ Greater chirality is associated with tighter folded α-helices.^21,30^

BLI demonstrated no significant differences between the BMP-2-affibody dissociation constants displayed by the recombinant and synthetic affibodies, indicating that these affibodies functioned similarly as binding proteins (Figure 1D). As expected, the synthetic high-affinity affibody displayed a significantly lower dissociation constant (i.e., higher BMP-2 binding affinity) than the synthetic low-affinity affibody.

### Synthetic affibodies contain fewer endotoxins than recombinant affibodies

Endotoxin content of the affibodies was measured before and after endotoxin removal via spin columns (Figure 2). The United States Food and Drug Administration (FDA) permits up to 0.5 EU/mL in medical devices.^7^ The recombinant affibodies contained more than the permissible amount of endotoxin, before and after treatment with the endotoxin removal columns. The endotoxin levels of the recombinant affibodies were significantly higher than those in synthetic affibodies, which were below the levels permissible by the FDA. These results were as expected since the recombinant affibodies were in close contact with bacterial lysate during purification, while the synthetic affibodies had no exposure to bacteria. Furthermore, endotoxins have high affinities towards metals such as nickel and cobalt which are used in protein collection, as well as for histidine tags which are present both during protein collection and during endotoxin removal, posing additional challenges for purifying recombinant proteins. ^28^

**Figure 2.**
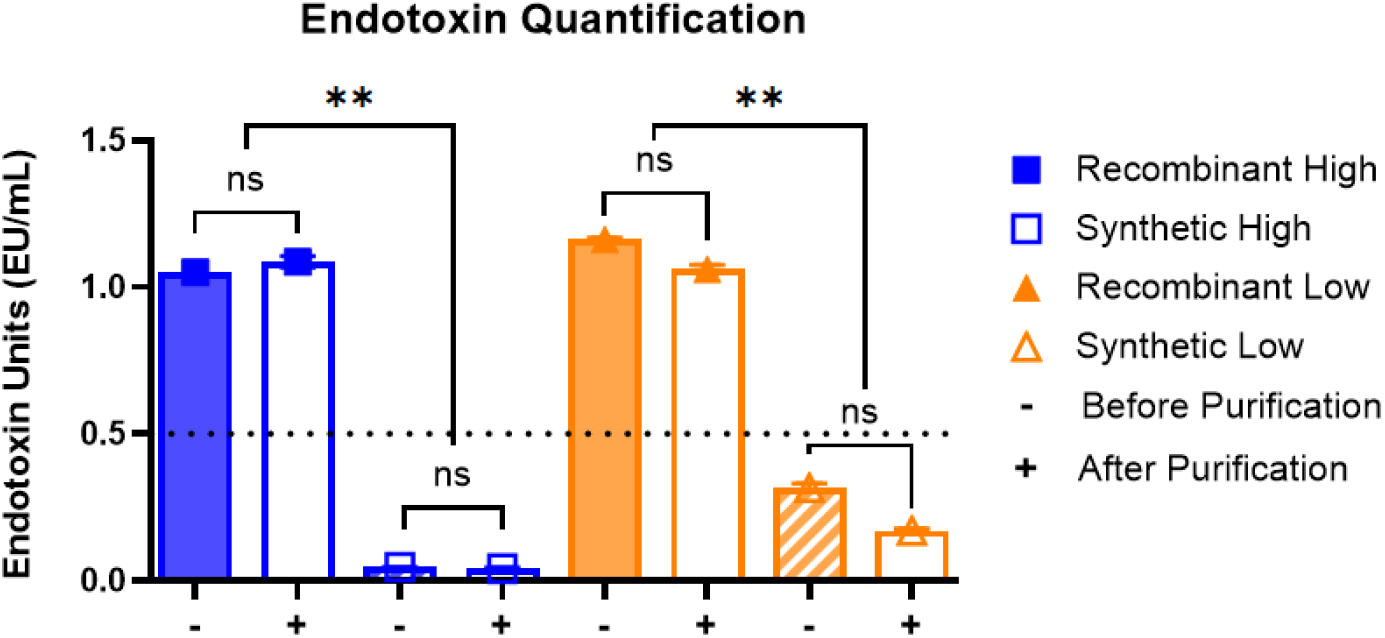
Endotoxin levels of recombinant and synthetic affibodies before and after endotoxin removal using spin columns. Endotoxin levels were measured using a chromogenic LAL endotoxin assay. Dotted line indicates the FDA approved limit for endotoxins in medical devices of 0.5 EU/mL. Statistical significance was determined using two-way ANOVA with Tukey post-hoc test. n=3; ns – not significant, ** p<0.01as indicated.

### Recombinant and synthetic affibodies similarly control BMP-2 encapsulation and release

PEG-mal hydrogels were synthesized without affibodies or with recombinant high-affinity, recombinant low-affinity, synthetic high-affinity, or synthetic low-affinity affibodies for BMP-2 encapsulation and release. Hydrogels containing high-affinity affibodies demonstrated a higher BMP-2 encapsulation efficiency (Figure 3A) and lower cumulative BMP-2 release (Figure 3B) than the PEG and low-affinity affibody hydrogels. There were no significant differences in BMP-2 encapsulation and release between recombinant and synthetic affibodies for either the high-affinity or low-affinity affibodies. The slopes of Fickian diffusion determined using the linear portion of each hydrogel release profile were significantly lower for both the recombinant and synthetic high-affinity affibody-conjugated hydrogels compared to the low-affinity and affibody-free hydrogels, indicating a lower effective BMP-2 diffusivity in these systems. No significant differences between the synthetic and recombinant affibodies were observed in Fickian diffusion slopes (Figure 3C). These results were consistent with our previously published work using recombinantly expressed affibodies.^5^

**Figure 3.**
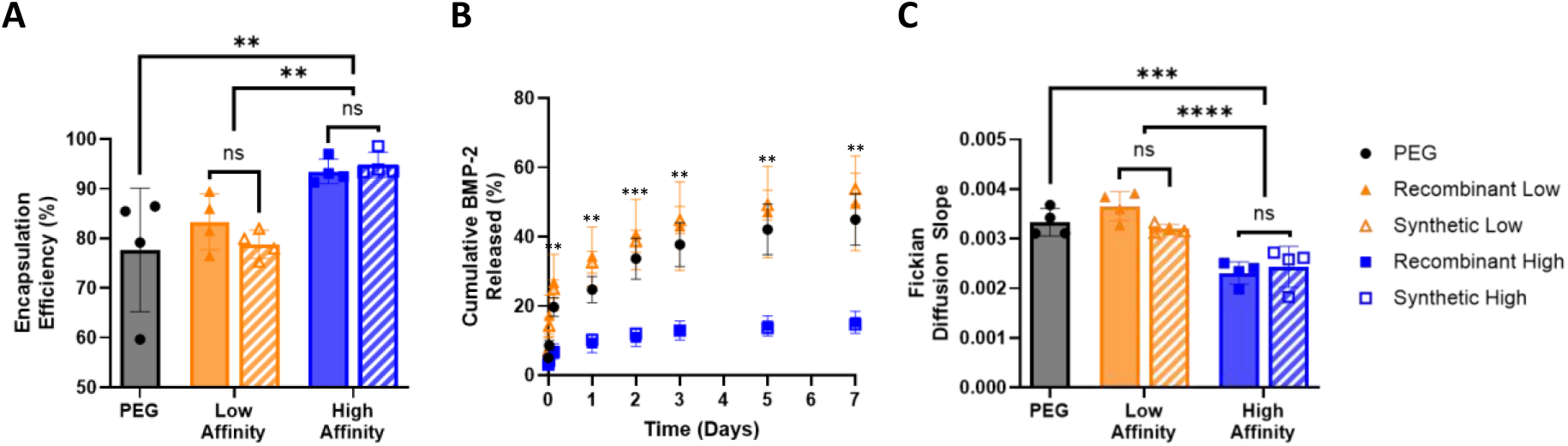
Encapsulation and cumulative release of BMP-2 from affibody-conjugated hydrogels. A) High-affinity affibody-conjugated hydrogels encapsulated more BMP-2 than low-affinity affibody and affibody-free hydrogels. B) High-affinity affibody-conjugated hydrogels released significantly less BMP-2 compared to low-affinity affibody and affibody-free hydrogels. C) Fickian diffusion rates. High-affinity affibody-conjugated hydrogels had a significantly slower Fickian diffusion rate compared to low-affinity and affibody-free hydrogels. Statistical significance was determined using 2-way ANOVA with Tukey post-hoc test. n=4; ns – not significant, ** p<0.01, *** p<0.001, **** p<0.0001; for panel B asterisks represent significance between the the high-affinity formulations and the PEG and low-affinity formulations.

## Discussion

Affibodies are versatile affinity-binding proteins that can be produced using recombinant protein expression or SPPS and can be used for various biomedical applications.^3,31–33^ Recent advances have enabled recombinantly expressed affibodies for controlled protein delivery using hydrogels.^2,4,5^ Here, we demonstrated that BMP-2-specific affibodies produced via SPPS have comparable structure and function to affibodies expressed recombinantly in *E. coli*.

Recombinant and synthetic affibodies share characteristic α-helical folding, similar dissociation constants for affibody-BMP-2 binding, and the ability to retain and release BMP-2 from PEG-mal hydrogels. Notable differences include lower molecular weights for the synthetic affibodies, which were engineered without hexahistidine tags, and could be leveraged for higher degrees of hydrogel modification, different magnitudes in circular dichroism signal, which could be representative of greater chirality in the synthetic affibodies, and lower endotoxin levels in the synthetic affibodies compared to their recombinant equivalents. These data suggest that synthetic and recombinant affibodies behave similarly in controlled BMP-2 release applications.

Although affibodies are typically expressed recombinantly in *E. coli*, SPPS may provide several advantages over recombinant protein expression. As noted in this work, recombinant expression of affibodies through an *E. coli* expression system yielded high levels of endotoxin contamination. Even after endotoxin removal, recombinant affibodies contained significantly higher endotoxin levels than synthetic affibodies, motivating the use of SPPS for affibody production for applications that eventually require in vivo use. Furthermore, SPPS has a high product yield and eliminates the risk of introducing other contaminating bacterial proteins into the product. Conversely, generating synthetic affibodies requires extensive infrastructure for SPPS that may not be readily available to all research groups. Generally, larger products also become more challenging to prepare by SPPS due to the stepwise growth of each product by chemical reaction. In certain cases, it may be advantageous to initially screen several affibodies generated recombinantly before using SPPS to make large quantities of a single affibody. Ultimately, this work reveals that recombinant and synthetic affibodies share comparable structure and function, demonstrating that these methods could be used interchangeably depending on the application and reaffirming the robust nature of these affibody molecules.

